# Venlafaxine stimulates an MMP-9-dependent increase in excitatory/inhibitory balance in a stress model of depression

**DOI:** 10.1101/794628

**Authors:** Seham Alaiyed, Mondona McCann, Gouri Mahajan, Grazyna Rajkowska, Craig A. Stockmeier, Kenneth J. Kellar, Jian Young Wu, Katherine Conant

## Abstract

Emerging evidence suggests that there is a reduction in overall cortical excitatory to inhibitory balance in major depressive disorder (MDD), which afflicts approximately 14-20% of individuals. Reduced pyramidal cell arborization occurs with stress and MDD, and may diminish excitatory neurotransmission. Enhanced deposition of perineuronal net (PNN) components also occurs with stress. Since parvalbumin-expressing interneurons are the predominant cell population that is enveloped by PNNs, which enhance their ability to release GABA, excess PNN deposition likely increases pyramidal cell inhibition. In the present study we investigate the potential for matrix metalloprotease-9 (MMP-9), an endopeptidase secreted in response to neuronal activity, to contribute to the antidepressant efficacy of venlafaxine, a serotonin/norepinephrine reuptake inhibitor. Chronic venlafaxine increases MMP-9 levels in murine cortex, and increases both pyramidal cell arborization and PSD-95 expression in the cortex of wild-type but not MMP-9 null mice. We have previously shown that venlafaxine reduces PNN deposition and increases the power of *ex vivo* gamma oscillations in conventionally-housed mice. Gamma power is increased with pyramidal cell disinhibition and with remission from MDD. Herein we observe that PNN expression is increased in a corticosterone-induced stress model of disease and reduced by venlafaxine. As compared to mice that receive concurrent venlafaxine, corticosterone treated mice also display reduced *ex vivo* gamma power and impaired working memory. Autopsy-derived prefrontal cortex samples show elevated MMP-9 levels in anti-depressant treated MDD patients as compared to controls. These preclinical and postmortem findings highlight a link between extracellular matrix regulation and MDD.

## Introduction

Major depressive disorder (MDD) is a common psychiatric illness afflicting 7-12% of men and 20-25% of women (Belmaker and Agam, 2008). Untreated MDD increases one’s risk for substance abuse, dementia, and suicide. Emerging evidence suggests that MDD is also associated with a reduction in overall excitatory to inhibitory (E/I) balance [reviewed in (Thompson et al., 2015; Page and Coutellier, 2019)]. For example, depressed patients show hypoactivity in the prefrontal cortex (PFC) (Fales et al., 2009). Similarly, animal models related to depression show atrophy of distal pyramidal dendrites in the PFC, loss of dendritic spines, and reduced expression of the AMPA glutamate receptor subunit GluA1 (Liston et al., 2006; Radley et al., 2008). In the hippocampus, chronically stressed animals also show atrophy of distal apical dendritic branches in excitatory neurons, and decreases in dendritic spine size and number (Magarinos and McEwen, 1995; Pawlak et al., 2005). A layer-specific decrease in GluA1 and post-synaptic density-95 (PSD-95) expression has also been described (Kallarackal et al., 2013). A better understanding of how E/I balance is altered in depression and how it may be restored by therapy could lead to the development of improved therapeutics.

Levels of matrix metalloproteases (MMPs), a family of zinc-dependent endopeptidases that play a role in adaptive and maladaptive learning and memory (Nagy et al., 2006; Meighan et al., 2007; Conant et al., 2010; Smith et al., 2014), may be increased by commonly prescribed monoamine reuptake inhibitors and their activity may also enhance E/I balance (Alaiyed et al., 2019). For example, monoamines stimulate G-protein coupled receptors that increase MMP expression, including the D1 dopamine receptor and the 5-HT-7 serotonin receptor (Li et al., 2016; Bijata et al., 2017). In terms of E/I balance, MMPs have been shown to stimulate dendritic arborization of pyramidal neurons as well as dendritic spine formation and expansion (Wang et al., 2008; Allen et al., 2016). In addition, through cleavage of perineuronal net components, MMPs may reduce glutamate input and membrane capacitance of parvalbumin-expressing GABA-releasing interneurons thereby disinhibiting pyramidal neurons to indirectly enhance E/I balance (Frischknecht et al., 2009; Tewari et al., 2018).

Herein we evaluate the possibility that MMP-9, which is released from neurons in a neuronal activity-dependent manner and is critical for learning and memory in varied brain regions (Nagy et al., 2006; Ganguly et al., 2013; Smith et al., 2014), mediates antidepressant-associated increases in E/I balance. We test this in both conventionally housed C57BL/6J mice and C57BL/6J mice subjected to a corticosterone-induced stress model of depression (David et al., 2009). The corticosterone model induces an anxiety and depressive phenotype in C57BL/6 mice, a strain that is resistant to several stress protocols (David et al., 2009). Corticosterone-induced phenotypic changes in C57BL/6 mice include abnormalities in novelty suppressed feeding and open field performance (David et al., 2009). Our focus is MMP-dependent effects on dendritic arbor in excitatory pyramidal neurons, as well as PSD-95 levels in hippocampal lysates, both of which have the potential to influence E/I balance. We also examine effectors of E/I balance that instead involve MMP-dependent effects on inhibitory GABA-releasing interneurons. These include the expression of PNN components, the power of gamma oscillations, the abundance of *ex vivo* sharp wave ripples (SWRs), and working memory performance. The power of gamma oscillations contributes to working memory capacity and is reduced in MDD (Yamamoto et al., 2014; Arikan et al., 2018). Moreover, gamma power is normalized with symptomatic remission (Arikan et al., 2018). To further explore the relevance of our findings to MDD, we also examine MMP-9 and tissue inhibitor of metalloproteinase-1 levels in postmortem PFC samples from control, MDD and antidepressant-treated subjects with MDD. In all studies, we focus on the hippocampus and/or prefrontal cortex, two regions implicated in MDD (MacQueen and Frodl, 2011).

## Materials and Methods

### I. Chemicals and reagents

(±) -Isoproterenol hydrochloride was purchased from U.S Pharmacopeia (USP catalogue number 35100). (±) -Venlafaxine and carbamoylcholine chloride (carbachol) were purchased Sigma Chemical (St. Louis, MO, USA; catalogue numbers PHR1736 and C4382, respectively). (±) -Norepinephrine (+)-bitartrate salt was purchased from Millipore-Sigma (catalogue number A0937), and sotalol hydrochloride was purchased from Tocris (catalogue number 0952).

### II. Cell culture and treatments

Primary hippocampal neuronal cells were prepared from WT post-natal day 0-1 mouse pups. The hippocampi were dissected after the removal of the meninges and dissociated into a single cell suspension with titration following 15 minutes incubation with 0.25% trypsin (Beaudoin et al., 2012). Neuronal cells were plated on 18 mm sterilized glass coverslips in a 12 well plate at approximately 90,000 cells concentration. Cultured cells were maintained in neural basal media supplemented with B27, glutamine, and penicillin/streptomycin antibiotic and incubated at 37 °C in a humidified 5% CO_2_-containing atmosphere for 18 days *in vitro* (DIV). Treatment was started at DIV 14-18. Cells were treated with isoproterenol (30 uM), norepinephrine (25 uM), or sotalol (20 uM) for 24 hours. Neuronal culture supernatants were concentrated 10 fold (VWR filter concentrators, catalogue number 82031-344) and saved for ELISA.

### III. Mice

Animals used for experiments were age-matched males and females, approximately 4 weeks old at the start of treatment. Strains were C57BL/6J (Jackson Laboratory, RRID: IMSR_JAX:000664) or MMP-9 homozygous null mice (Jackson Laboratory, RRID: IMSR_JAX:007084) that had been backcrossed to C57BL/6J mice for at least five generations. Experimental groups were matched in terms of the male to female ratio for each comparison, and then arbitrarily assigned to treatment groups. Mice were housed four-five per cage. Food and water were provided *ad libitum*. Experiments were performed in accordance with National Institutes of Health guidelines and institutionally approved protocols (2016-1117 and 2018-0037). Cages were supplied with balconies and nesting materials for enrichment.

### IV. Corticosterone and venlafaxine treatment

Mice that received saline or venlafaxine (VFX) were treated for two weeks with a daily IP injection of sterile saline or 30mg/kg VFX in sterile saline (200 μl total volume). For the depression model used herein (David et al., 2009), mice were treated with corticosterone (35 ug/ml equivalent to 5 mg/kg/day) dissolved in 0.45 % β-cyclodextrin (β-CD) for 6 weeks and added to their drinking water. Control mice received 0.45 % β-CD in their drinking water. For experiments that also evaluated the effects of VFX in this model, during the last 2 weeks of the protocol mice were assigned to groups in which they received intraperitoneal VFX (30 mg/kg) or an equal volume of saline daily. To minimize animal distress, injections were delivered by trained veterinary technicians and the delivery site alternated from right to left each day.

### V. T maze and elevated plus maze (EPM) testing

EPM testing was performed using a standard apparatus and an ANY-maze video tracking as previously described (Allen et al., 2016). Working memory was assessed by T maze testing (Shoji et al., 2012). The T shaped apparatus was placed with the stem of the T proximal to the experimenter. Testing was begun with the distal right and left drop down doors raised. Unlike the reward version of this test, no habituation to the maze is used, as it is the novelty of the maze that tests spontaneous alternation. Each mouse was placed in the start arm with the door lowered, and following removal of this door the animal was allowed to freely choose to enter either the right or left arm. After the mouse chose a goal arm, the door was lowered to confine the mouse in that arm for 30 seconds. The mouse was then removed gently, placed into a rest cage from which the maze was not visible, the maze was cleaned, and the drop down doors to the distal right and left arms raised. At that point the same mouse was again placed in the start arm and the test repeated. This was done sequentially for each mouse in the cohort, and then sequentially repeated. Choices were recorded for each trial as a 1 (alternation) or 0 (no alternation).

### VI. Sharp wave ripple and gamma recordings

Hippocampal slices and local field potential (LFP) recordings of gamma-frequency oscillations and sharp wave ripples (SWR) were prepared as previously described (Alaiyed et al., 2019). Briefly, following the last day of VFX or saline injection mice were anesthetized by isoflurane inhalation. After confirming unconsciousness by an inability to respond to a strong paw pinching, animals were rapidly decapitated and the whole brain extracted and chilled in cold (0°C) sucrose-based cutting artificial cerebrospinal fluid containing (in mM) 252 sucrose; 3 KCL; 2 CaCl2; 2 MgSO4; 1.25 NaH2PO4; 26 NaHCO3; 10 dextrose and bubbled with 95% O_2_, 5% CO_2_. Hippocampal slices (490 μm thick) were cut in horizontal sections from dorsal to ventral locations with a vibratome (Leica, VT1000S). To minimize potential dorsal to ventral activity differences, we recorded from the 6-8^th^ slices obtained when cutting began at the dorsal most aspect of the brain. Slices were transferred into a chamber filled with ACSF contained (in mM) NaCl, 132; KCl, 3; CaCl2, 2; MgSO4, 2; NaH2PO4, 1.25; NaHCO3, 26; dextrose, 10; and saturated with 95% O2 and 5% CO2 at 26°C. Slices were incubated for at least 180 min before being moved to the recording chamber. For LFP recordings of gamma-frequency oscillations and SWR, low-resistance glass microelectrodes (approximately 150K tip resistance) were used. Electrodes were filled with 1 M NaCl. Recordings were performed in CA1 stratum oriens proximal to CA2. The ACSF fluid was switched to ACSF containing carbachol at 40 µM for recording of carbachol-stimulated gamma oscillations. Local field analyses were performed with a previously described custom MATLAB algorithm, available upon request (Alaiyed et al., 2019).

### VII. Preparation of brain lysates and fixed tissue

Following treatment and euthanasia, brains were bisected into right and left hemi brains. Homogenates from hippocampal and cortical tissue were prepared by lysis in immunoprecipitate buffer [50 mM Tris, pH 7.5, 150 mM NaCl, 0.1% sodium dodecyl sulfate, 1% octylphenoxypoly (ethyleneoxy) ethanol, branched, and 1X protease and phosphatase cocktail (Thermo Scientific 1861281)]. Lysates were sonicated for 10 seconds, placed on ice for 20 minutes, and centrifuged 15 minutes at 14,000 rpm at 4 °C. Lysate supernatants were saved for protein analyses. For immunohistochemistry, hemi-brains were rapidly fixed in chilled 4% PFA overnight at 4 °C prior to paraffin embedding and thin slice preparation (15 μm).

### VIII. ELISA and Western blot

Pro MMP9 protein concentrations in cell culture supernatants and tissue lysates were measured by ELISA, performed according to the manufacturer’s instructions (Mouse Pro-MMP-9, R&D systems catalogue number MMP900B). For immunoblotting, protein concentrations were determined using a BCA protein assay (Pierce Biotechnology, Inc.), and equal amounts of protein were loaded in each lane. Samples (50 µg total protein) were immersed in Laemmli sample buffer (Bio-Rad, Hercules, CA, USA, catalog #161-0737) containing 5% β-mercaptoethanol, and boiled for 5 minutes at 95 °C. Samples were subsequently separated by electrophoresis on precast gels (4-20% mini protein TGX gels, Bio-Rad catalog #456-1094) and transferred to nitrocellulose membranes (Trans-Blot Turbo Transfer, Bio-Rad, catalog #1704159). Membranes were probed with primary and secondary antibodies, and bands visualized by chemiluminescence as previously described (Alaiyed et al., 2019). Primary antibodies used were mouse anti-Brevican (1:1000, 610894, BD Transduction Laboratories), mouse anti-PSD-95 (1:1000, CP35, Millipore), and mouse anti-GAPDH (1:5000, MAB374, Millipore).

### IX. Golgi staining and analysis

Golgi staining was performed with the FD Rapid Golgi Stain Kit (FD NeuroTechnologies, INC catalog #PK401) according to the manufacturer’s instructions. After euthanasia, murine brains were quickly removed, rinsed in double distilled water and immersed in the impregnation solution. Brains were stored in this solution at room temperature for 2 weeks in the dark. Tissue was subsequently transferred to solution C and stored at room temperature for 1 week. Brains were sectioned (150 μm) with a vibratome (Leica, VT1000S). Each section was then transferred and mounted on gelatin-coated microscope slides and allowed to dry naturally at room temperature in the dark. After rinsing in double distilled water 2 times, 4 minutes each, slides were put into the manufacturer-recommended mixture of staining solutions for 10 minutes then rinsed. Sections were dehydrated in 50%, 75%, 95%, 100% ethanol, and tissue-clear for 4 minutes each and then mounted with Hydromount (National Diagnostics, HS-106). Visualization of arbors and quantification was subsequently performed as described (Allen et al., 2016). Four to five mice were used per group, and 1-2 slides were analyzed per mouse (total 9-10 slides/mouse). Slide identity was coded and quantification of arbors was performed by an investigator blind to cohort.

### X. Immunostaining

Immunostaining was performed as described (Alaiyed et al., 2019). Briefly, following fixation in 4% PFA, brains were paraffin-embedded and sectioned (15 microns). Sections were washed 2-3 times with 1X-PBS, then permeabilized with 1X-PBS containing 0.1% Triton X-100, blocked with 10% normal goat serum, and incubated with anti-parvalbumin (1:500, Sigma, P3088) overnight at 4 °C. Following subsequent washes and incubation with a fluorescent secondary antibody for PV immunostaining and fluorescein-labeled Wisteria floribunda lectin (WFA) (1:1000, Vector Laboratories, FL-1351) for 2 hours at room temperature, sections were washed several times with 1X-PBS, counterstained with DAPI and mounted with Hydromount (National Diagnostics, HS-106) before being allowed to dry several days at 4 °C. For PV and PNN cell quantification, images were acquired using a Leica SP8 laser scanning confocal microscope with an oil immersion, 209 objective with .40 numerical aperture. Laser intensity, gain, and pinhole settings were kept constant for all samples. Images were taken through a z-plane (8.5 lm) within the center of the tissue section, containing 20 stacks (0.4 lm/stack) from the dorsal hippocampus. Quantification of PV numbers with and without an associated PNN was counted for each image that was acquired from regions of interest (ROI) using 8-10 mice per group, and 2-3 slides per mouse. For PNN intensity, a semi-automated analysis, ‘PIPSQUEAK’ macro plugin, was used [described in (Slaker et al.2016). This macro uses an ROI protocol for double-labeled cell-based quantification. Following background subtraction within an appropriate radius, WFA or PV cells are identified based on threshold limit requirements. ROIs were identified around WFA-labeled PV-expressing cells and the mean intensity was measured. Data points represent averages for ROIs from one to two images from each of animal.

### XI. Human tissue analyses

The Institutional Review Boards of the University of Mississippi Medical Center, Jackson, MS, and the University Hospitals Cleveland Medical Center, Cleveland, OH, approved all procedures of tissue collection and retrospective psychiatric assessment, as previously described (Ho et al., 2019). Informed consent was obtained from legally-defined next-of-kin for tissue collection, interviews, and medical records. A master-level social worker administered the Structured Clinical Interview for DSM-IV Axis I Disorders to informants with knowledge of the subjects (First et al., 2002). A board-certified clinical psychologist and a board-certified psychiatrist independently reviewed the scoring of the diagnostic interview, medical records, and the medical examiner’s report. Psychopathology of the subjects, or the lack thereof in control subjects, was determined in consensus with the clinical psychologist, the psychiatrist, and the social worker. Subjects meeting DSM-IV criteria for MDD, or controls without a psychiatric diagnosis were selected for this study. Subjects with any neuropathological or neurological disorders were excluded. The medical examiner’s office assessed the presence of psychotropic medications and substances of abuse in blood and urine.

Samples of postmortem brain tissue were acquired at autopsy at the Cuyahoga County Medical Examiner’s Office, Cleveland, OH. The medical examiner determined the cause of death. Gray matter tissues from the frontal pole (Brodmann’s area 10) were dissected and rapidly frozen in 2-methylbutane cooled by dry ice, and subsequently buried in powdered dry ice prior to storage at −80 °C. Samples were segregated into three groups, age- and sex-matched, and coded so that research staff were unaware of diagnoses. The groups (n=12-15) included psychiatrically normal control subjects, subjects with MDD but had not been treated with an antidepressant medication in the weeks prior to death, and subjects with MDD that were treated with an antidepressant medication having a proximal effect on the monoamine neurotransmitter system. Antidepressant treatment was confirmed by postmortem toxicology in MDD subjects and these subjects also had an active prescription in the last month of life. The mean age, age range, and post mortem delay ± S.D. for each group was as follows: Control (age 51 ± 15.1, postmortem delay 22.2 h ± 7.5); MDD (age 50 ± 15.7, range 28-83; postmortem delay 22 h ± 6.5); MDD plus antidepressant (age 52 ± 16.5, postmortem delay 23.6 h ± 8.3). Brain lysates were prepared according to established methods (St Hillaire et al., 2005), and ELISA for MMP-9 and TIMP-1 was performed using commercially available kits (R and D Systems, Minneapolis, MN). Data were analyzed and corrected for total protein levels prior to sending it to co-investigators to break the diagnostic code on samples.

### XII. Experimental design and statistical analyses

Sample size was based on experience indicating that 4-8 animals per group are typically necessary to observe physiologically relevant changes in MMP levels and additional biochemical endpoints (Alaiyed et al., 2019). Experiments focused on PNN intensity and population recordings were set up to utilize 4-6 animals and several slices from each as indicated in the methods section. Data were entered into a Graph Pad Prism 8.0 program and statistical analysis was performed using Student’s unpaired t-test for two group comparisons or ANOVA, with post-hoc analyses as indicated, for comparisons of more than two groups. Multiple comparisons were controlled for and within- and between-subject comparisons were considered as indicated in the results for human subject data. Significance was set at *p* < 0.05 and ROUT testing was performed to identify outliers.

## Results

### 1. β adrenergic receptor activation increases MMP-9 release from cultured neurons and MMP-9 levels are increased in brain lysates from venlafaxine-treated animals

Previous studies have demonstrated that 5HT-7 serotonin receptor activation, which is linked to Gαs stimulation, increases MMP-9 activity in cultured neurons (Bijata et al., 2017). Higher doses of venlafaxine (VFX), as used in this study, can also increase extracellular levels of norepinephrine. We therefore examined agonists of Gαs coupled norepinephrine receptors for their ability to stimulate an increase in MMP-9 release from cultured murine hippocampal neurons. As shown in figure 1A, the non-selective β adrenergic agonist isoproterenol stimulated a significant increase in the release of MMP-9 from cultured murine hippocampal neurons (n= 5-6 biological replicates, *p*=0.0164, one-way ANOVA with Tukey’s post-hoc testing). While the difference between isoproterenol and norepinephrine was not significant, the difference between isoproterenol and all other groups did reach significance.

**Figure 1.**
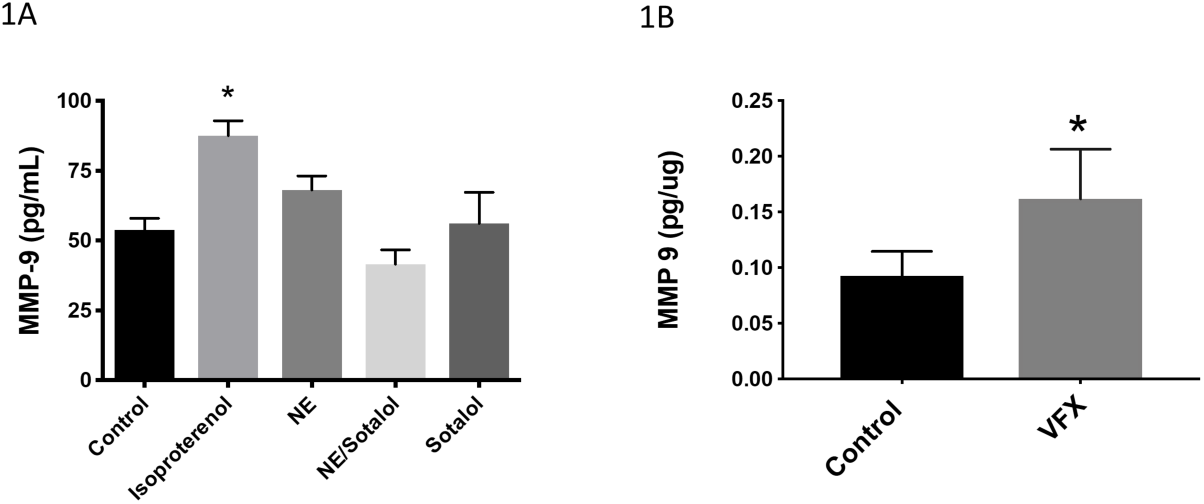
β adrenergic receptor activation increases MMP-9 release from cultured neurons and MMP-9 levels are increased in brain lysates from venlafaxine treated animals. As shown in figure 1A, the non-selective β adrenergic agonist isoproterenol stimulated a significant increase in the release of MMP-9 from cultures of murine hippocampal neurons (n= 5-6 biological replicates, *p*=0.0164, one-way ANOVA with Tukey’s post-hoc testing). In addition, while the difference between isoproterenol and norepinephrine was not significant, the difference between isoproterenol and all other groups did reach significance. Figure 1B shows MMP-9 levels in cortical lysates from animals that were treated for two weeks with saline or venlafaxine. Venlafaxine was associated with a significant increase in cortical MMP-9 (pg/μg total tissue, n=4-5 mice per group, *p*= 0.0328, Student’s t-test).

Because serotonin and norepinephrine reuptake inhibitors will indirectly activate monoamine receptor subtypes linked to varied downstream pathways, it was important to also assess MMP-9 levels in animals treated with monoamine reuptake inhibitors. We previously demonstrated VFX increases levels of MMP-9 in murine hippocampus (Alaiyed et al., 2019). Since some of the experiments in the present study are instead focused on the cortex, in figure 1B we examined MMP-9 in sensorimotor cortical lysates from animals that were treated for two weeks with saline or the VFX. Similar to what was observed in the hippocampus, VFX was associated with a significant increase in cortical expression of MMP-9 (pg/μg total tissue, n=4-5 mice per group, *p*= 0.0328, Student’s t-test).

### 2. VFX stimulates increased dendritic arbor and PSD-95 levels in wild type but not MMP-9 null mice

MMPs have been shown to enhance dendritic arborization and spine formation in varied settings (Wang et al., 2008; Verslegers et al., 2015; Allen et al., 2016). Potential mechanisms include their ability to activate pro-neurotrophins (Lee et al., 2001), their ability to generate integrin-binding ligands (Lonskaya et al., 2013), and their ability to modulate extracellular matrix in a manner that may be permissive for spine growth (Fawcett et al., 2019). To determine whether VFX had the potential to enhance dendritic arbor and spine formation in conventionally housed mice, we treated animals for 14 days with 30 mg/kg VFX and subsequently performed Golgi analyses to examine arborization and Western blots to quantify levels of PSD-95, which is typically increased in the setting of dendritic spine expansion and/or increased formation. Because the high density of dendritic arbors in hippocampus hinders quantification, arborization was quantified in the sensorimotor cortex. Representative images are shown in figure 2A and quantitation of branching data for primary, secondary and tertiary processes are shown in 2B-C. Results from PSD-95 quantification in hippocampal lysates are shown in 2D-G. As shown in figures 2B-C, VFX significantly increases the number of secondary dendrites in wild type but not MMP-9 null mice (n=39-45 neurons from 6-8 mice per group, wild type control versus VFX: *p*= 0.0176, Student’s t-test; MMP-9 null control versus VFX: not significant). And while tertiary dendritic branching appears increased by VFX in both the wild type and MMP-9 knockouts, the difference in tertiary branches for saline and VFX was not significant in either group, possibly due to a lower number of tertiary as compared to secondary processes. Note: normalized values are shown since the wild type +/- VFX and the MMP-9 null +/- VFX groups were stained at different times. As shown in 2D-G, VFX significantly increased PSD-95 levels in wild type mice (n= 6, *p*= 0.0388, Student’s t-test). There was, however, no significant difference in PSD-95 levels between the saline and VFX treated MMP-9 knockouts.

**Figure 2.**
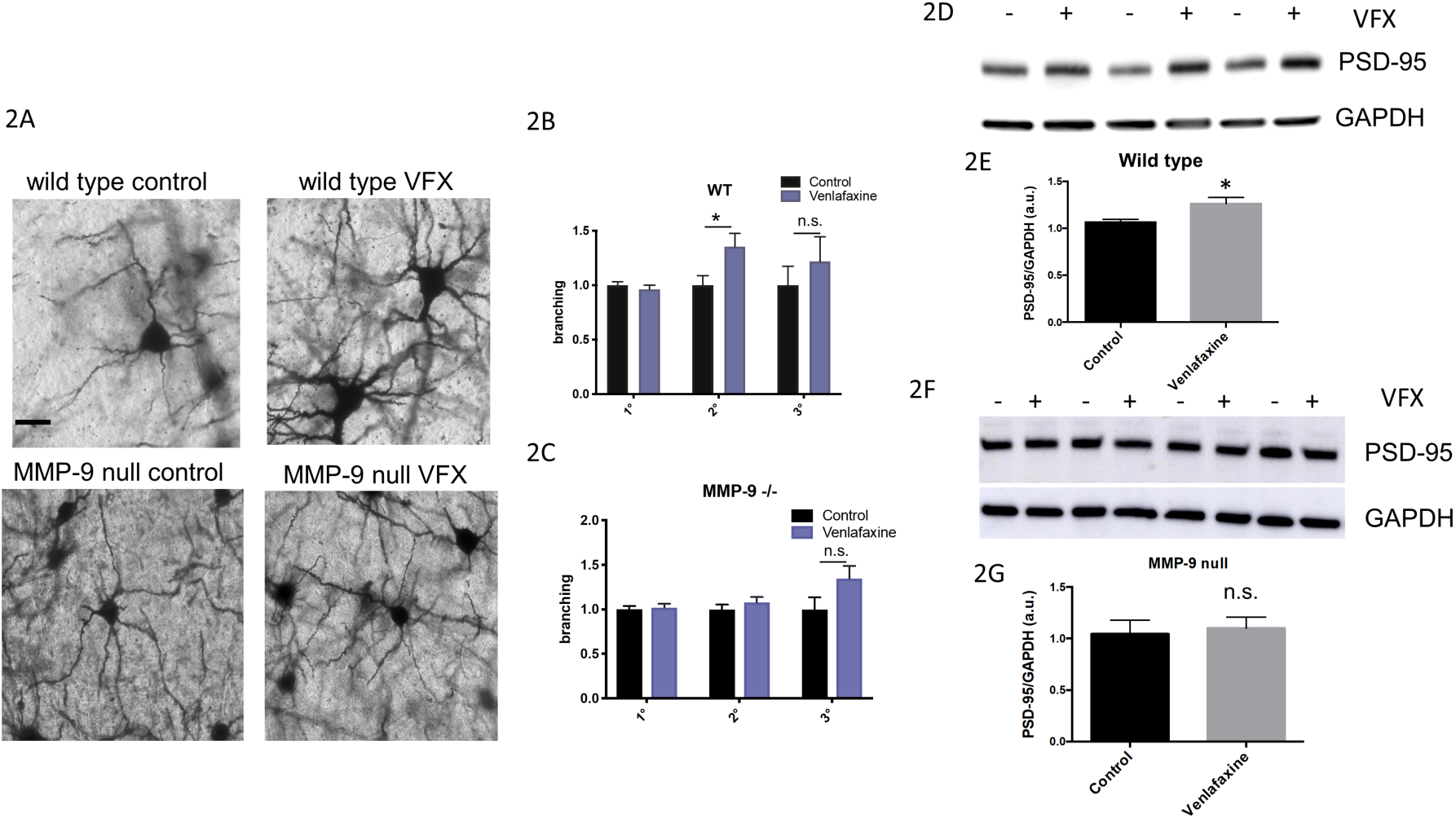
Venlafaxine stimulates increased dendritic arbor and PSD-95 levels in wild type but not MMP-9 null mice. Representative images from Golgi stained cortex for mouse strains and treatment groups are shown in figure 2A, and normalized branching data for primary, secondary and tertiary processes are shown in 2B-C. Results from PSD-95 quantification in hippocampal lysates are shown in 2D-G. As shown in figures 2B-C, VFX significantly increases the number of secondary dendrites in wild type but not MMP-9 null mice (n=39-45 neurons from 6-8 mice per group, wild type control versus VFX: *p*= 0.0176, Student’s t-test; MMP-9 null control versus VFX: not significant). As shown in 2D-G, VFX significantly increased PSD-95 levels in wild type mice (*p*= 0.0388, Student’s t-test). There was, however, no significant difference in PSD-95 levels between the saline and VFX treated MMP-9 knockouts. The scale bar represents 20 μm.

### 3. VFX increases brevican (BCAN) cleavage in wild type but not in MMP-9 null mice

While select changes in pyramidal cell structure can enhance E/I balance, reduced inhibitory input to pyramidal cells can do the same. PV-expressing interneurons are among the GABA releasing neuronal subtypes that can inhibit cortical and hippocampal pyramidal cells. PV expressing interneurons are also the predominant neuronal population that is surrounded by the PNN, a dense form of extracellular matrix that can be cleaved by MMPs. PNNs provide PV neurons with increased membrane capacitance and also localize glutamate input to synaptic contacts, thus facilitating the release of GABA from PV interneurons and thereby inhibiting pyramidal neuron activity (Frischknecht et al., 2009; Tewari et al., 2018). Disruption of the PNN can instead reduce PV-mediated pyramidal cell inhibition (Hayani et al., 2018).

To examine the potential for VFX to upregulate cleavage of specific PNN components, we examined the integrity of BCAN in saline and VFX treated animals. We also examined basal and VFX-stimulated BCAN levels and cleavage in wild type as compared to MMP-9 null animals. We focused on BCAN because it is a critical PNN component with respect to PV interneuron function (Favuzzi et al., 2017). In figures 3A-3C, we show full length and cleaved BCAN (80 kDa) in control and MMP-9 null mice. Representative Western blot results are shown in 3A and densitometric results from 4-5 mice per group are shown in 3B and 3C. It can be appreciated that while full length BCAN does not significantly differ between wild type and MMP-9 null mice (*p*= 0.24, Students t-test), basal BCAN cleavage is significantly reduced in MMP-9 null animals (*p*= 0.0004, Student’s t-test). In figures 3D-F we show full length and cleaved BCAN in saline and VFX treated wild type and MMP-9 knockout mice. Representative Western blot results are shown in 3D and 3E. Densitometric analyses with subsequent quantification of the ratio of cleaved/full length BCAN from n=4-5 mice per group are shown in figure 3F. The difference between control and VFX in wild type mice is significant (*p*=0.0278, 2-way ANOVA with Sidak’s multiple comparisons). In contrast, the difference between control and VFX in MMP-9 null animals is not significant (*p*>0.05, 2-way ANOVA with Sidak’s multiple comparisons).

**3.**
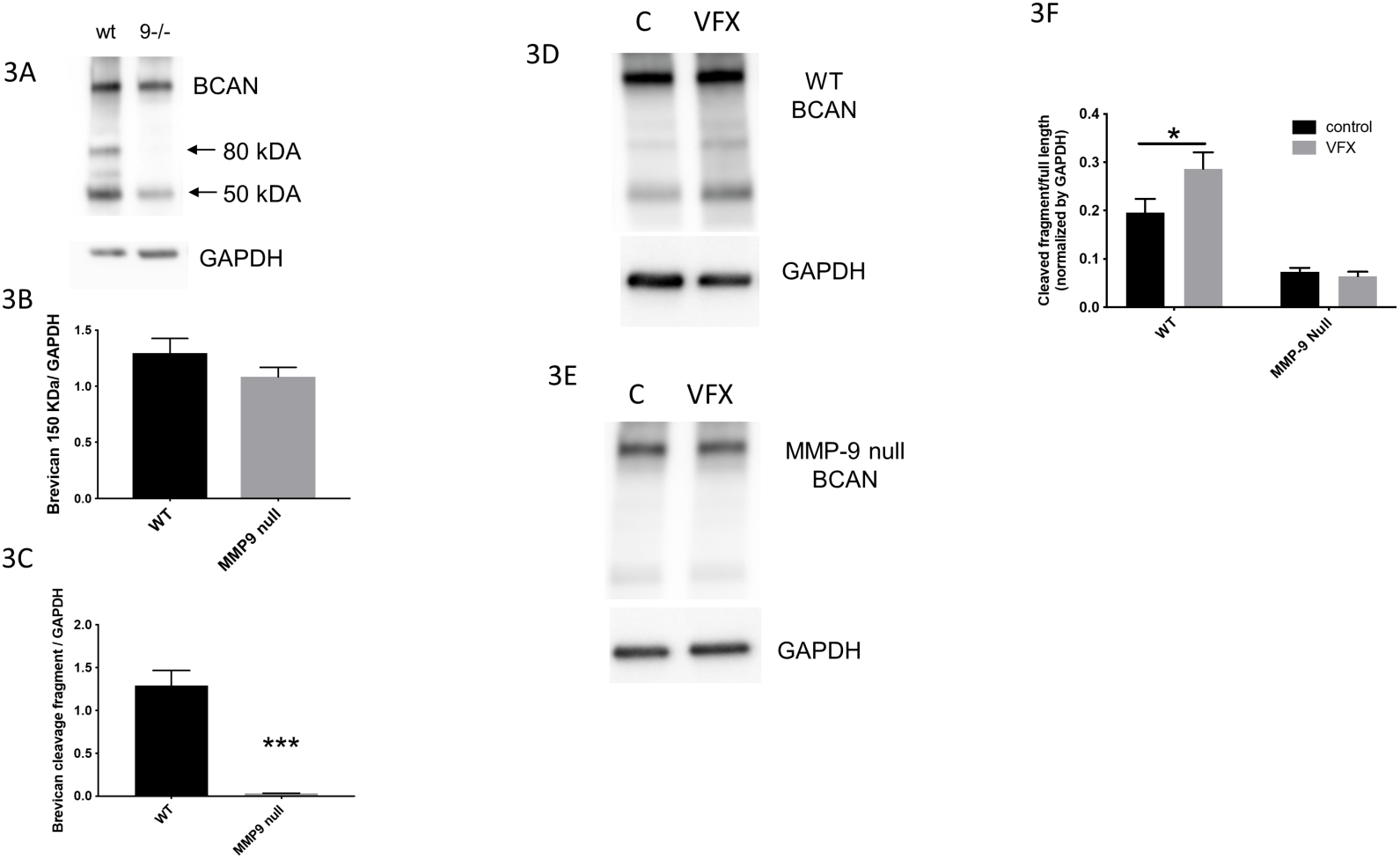
VFX increases brevican cleavage in wild type but not in MMP-9 null mice. Figures 3A-3C show full length and cleaved BCAN (80 kDa) in control and MMP-9 null mice. Representative Western blot results are shown in 3A and densitometric results from 4-5 mice per group are shown in 3B and 3C. It can be appreciated that while full length BCAN does not significantly differ between wild type and MMP-9 null mice (*p*= 0.24, Students t-test), basal BCAN cleavage is significantly reduced in MMP-9 null animals (p 0.0004, Student’s t-test). In figures 3D-F we show full length and cleaved BCAN in saline and VFX treated wild type and MMP-9 knockout mice. Representative Western blot results are shown in 3D and 3E. Densitometric analyses with subsequent quantification of the ratio of cleaved/full length BCAN from n=4-5 mice per group are shown in figure 3F. The difference between control and VFX in wild type mice is significant (*p*=0.0278, 2-way ANOVA with Sidak’s multiple comparisons). In contrast, the difference between control and VFX in MMP-9 null animals is not significant (*p*>0.05, 2-way ANOVA with Sidak’s multiple comparisons).

### 4. Corticosterone (CORT) increases BCAN and PNN levels

While emerging evidence links stress and depression to a reduction in E/I balance, most studies have focused on changes in pyramidal cell structure and function that would be expected to diminish the ability of pyramidal cells to respond to glutamate. Few studies have explored the possibility that depression could be secondary to increased GABA mediated inhibition. Since enhanced deposition of PNN components has the potential to increase PV interneuron-mediated inhibition, we examined BCAN levels in mice exposed to a corticosterone-induced stress model of depression. We focused on BCAN expression in the hippocampus as this is a region rich in receptors for CORT (Anderson et al., 2006). As shown in figure 4A-B, BCAN levels are increased in hippocampal lysates from CORT-exposed mice as compared to controls (n=4 mice per group, *p*=0.0466, Student’s t-test). Western blot data are shown in 4A and the results from densitometric analyses in 4B. Because previous results suggest that MMP-9 is an important mediator of basal BCAN cleavage, as well as and VFX-stimulated PNN cleavage (Alaiyed et al., 2019), we also looked at ratios of MMP-9 and its inhibitor, TIMP-1, in hippocampal lysates from CORT-exposed animals treated with saline or VFX. As shown in figure 4C, VFX significantly increased MMP-9/TIMP-1 levels over control (n=5-6 mice per group, *p*= 0.0027, one-way ANOVA with Tukey’s multiple comparisons) while CORT alone had no significant effect as compared to control (*p*= 0.094, one-way ANOVA with Tukey’s multiple comparisons).

**4.**
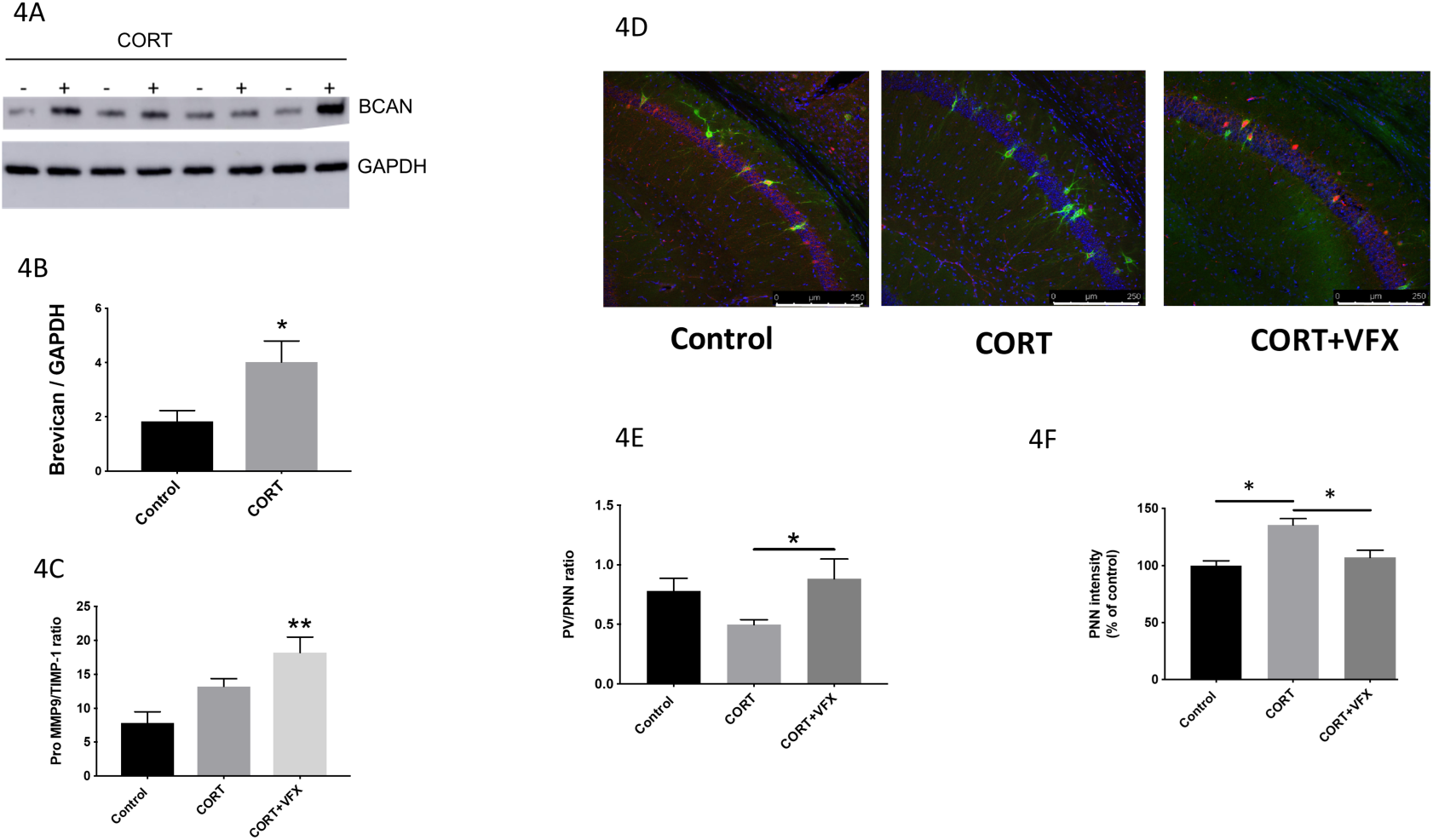
Corticosterone (CORT) increases BCAN and PNN levels. BCAN levels in hippocampal lysates from control and CORT-exposed mice are shown in Figures 4A-B (Western blot and densitometry respectively). BCAN levels are increased in hippocampal lysates from CORT-exposed mice as compared to controls (n=4 mice per group, *p*=0.0466, Student’s t-test). Ratios of MMP-9 and its inhibitor, TIMP-1, in hippocampal lysates from CORT-exposed animals treated with saline or VFX are shown in Figure 4C. VFX significantly increased MMP-9/TIMP-1 levels over control (n=5-6 mice per group, *p*= 0.0027, one-way ANOVA with Tukey’s multiple comparisons) while CORT alone had no significant effect as compared to control (*p*= 0.094, one-way ANOVA with Tukey’s multiple comparisons). Figure 4D shows representative PNN staining for control, CRT and CORT +VFX exposed wild type animals, while figures 4E-F we show quantification of the PV/PNN ratio and PNN intensity. VFX significantly increases the PNN/PV ratio in CORT-exposed animals (6-8 mice, n = 18-26 slides, *p*= 0.0383, ANOVA with Tukey’s multiple comparisons) and reduces overall PNN fluorescence intensity (*p*< 0.0008), consistent with a reduction in PNN levels. The scale bar represents 250 μm.

We have previously shown that VFX stimulates an MMP-9 dependent decrease in PNN intensity in conventionally housed mice (Alaiyed et al., 2019). We here examined the effects of VFX on overall PNN levels in the CORT model. In figure 4D we show representative staining while in figures 4E-F we show quantification of the PV/PNN ratio and PNN intensity in wild type animals. VFX significantly increases the PNN/PV ratio in CORT-exposed animals (6-8 mice, n = 18-26 slides, *p*= 0.0383, ANOVA with Tukey’s multiple comparisons) and reduces overall PNN fluorescence intensity (*p*< 0.0008), consistent with a reduction in PNN levels.

### 5. VFX stimulates an MMP-9-dependent increase in the power of gamma oscillations in ex vivo hippocampal slices from CORT treated animals

Since chronic CORT increased BCAN levels, and because the PNN is thought to facilitate the firing of PV interneurons and thus pyramidal cell inhibition, we next examined the power of gamma oscillations. Gamma power is increased with PNN attenuation (Lensjo et al., 2017) and correlated with the magnitude of pyramidal cell mediated sharp waves. Importantly, gamma power is reduced in depression and animal models of depression (Khalid et al., 2016; Arikan et al., 2018). We previously demonstrated that VFX could increase carbachol-stimulated gamma power in hippocampal slices from conventionally housed, non-CORT-treated, animals (Alaiyed et al., 2019). Carbachol is used in slice studies of gamma to substitute for septal cholinergic input. As shown in figures 5A and B, VFX also increases low and high gamma power in CORT-treated animals. The difference between CORT and CORT +VFX groups is significant (4-6 mice, n= 10-13 slices, low gamma *p*=0.0004, high gamma *p*= 0.0026, one-way ANOVA with Sidak’s multiple comparisons) as is the difference between control and CORT in the low gamma range (p= 0.045, one-way ANOVA with Sidak’s multiple comparisons). In contrast the difference between controls and CORT +VFX is not significant. Shown in 5C are representative tracings from the 3 groups as indicated. We also evaluated the MMP-9 dependence of VFX’s effect on gamma power in the CORT model. As shown in figures 5D and E, there is no significance difference between the CORT and CORT +VFX groups in MMP-9 knockout animals (4-5 mice, n= 9-11 slices, low gamma *p*= 0.96 high gamma *p*=0.63, one-way ANOVA with Sidak’s multiple comparisons). In 5F we show representative tracings from the knockout animals. While the CORT and CORT + VFX groups show no difference in the MMP-9 knockouts, it also appears that CORT has a lesser effect on gamma in the MMP-9 nulls as compared to wild type animals (control versus CORT conditions). One possibility is that MMP-9 could play a role in both CORT and VFX associated changes in gamma. CORT-stimulated increases in MMP-9 might activate glia to enhance glial expression of PNN proteins, while VFX may increase MMP-9 in proximity to neurons to more than overcome effects of CORT in wild type animals. Future studies will be necessary to address this possibility and others.

**5.**
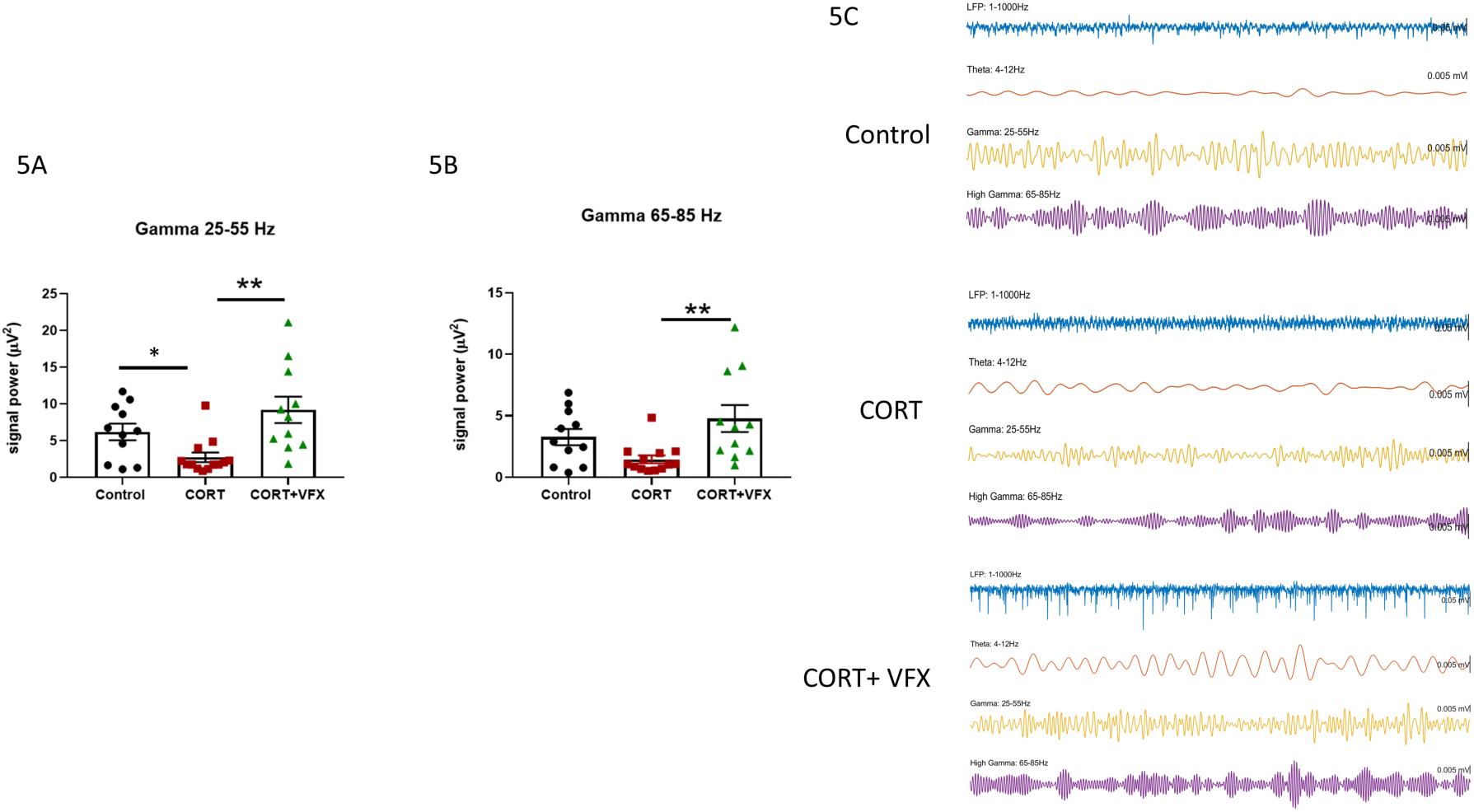
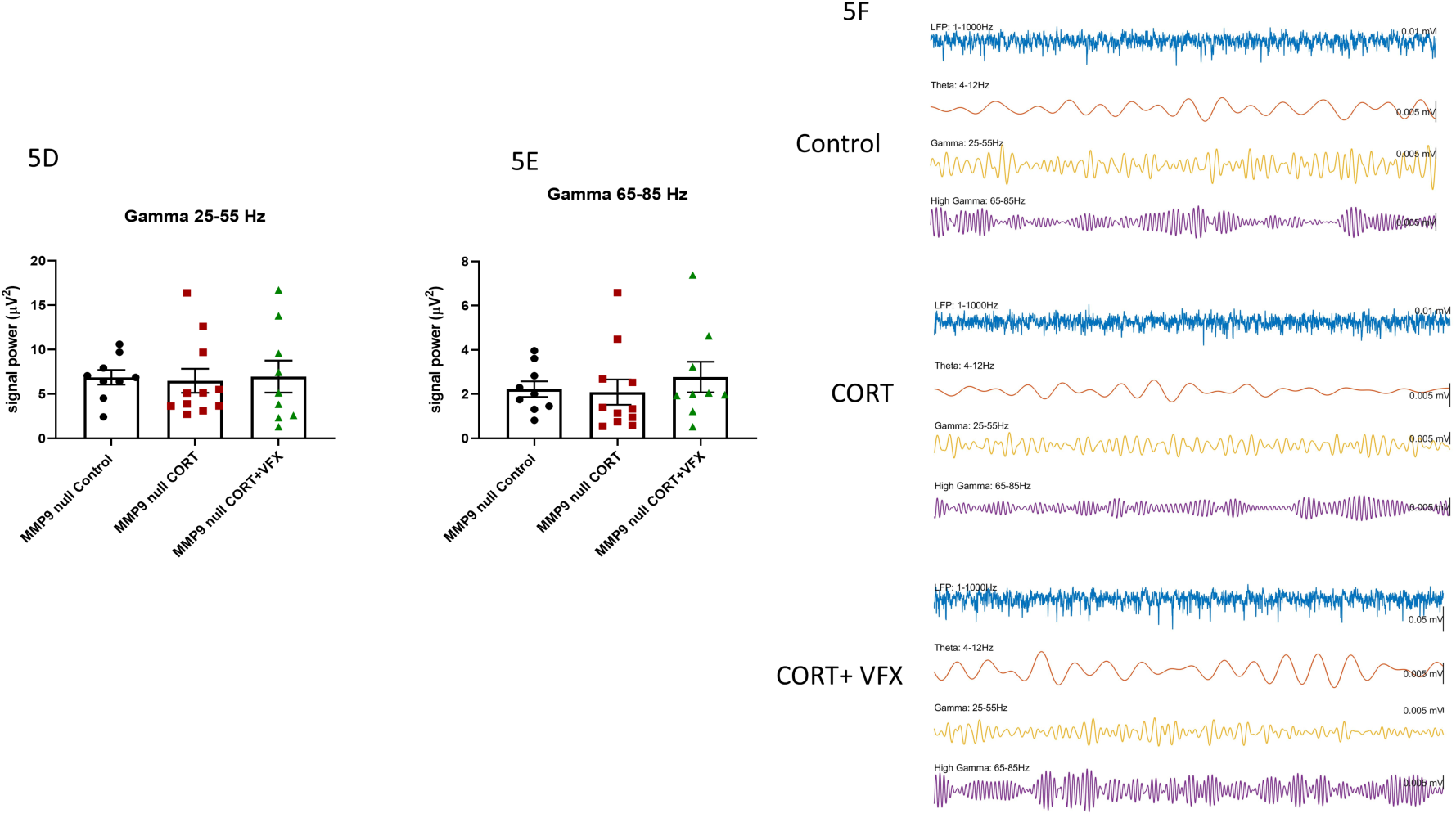
Venlafaxine stimulates an MMP-9-dependent increase in the power of gamma oscillations in ex vivo hippocampal slices from CORT treated animals. Shown in figures 5A and B, are results from local field potential recordings of carbachol – stimulated low and high gamma power in *ex vivo* hippocampal slices. As also shown in figure 5A and B, venlafaxine increases low and high gamma power in CORT-treated animals. The difference between CORT and CORT +VFX groups is significant (4-6 mice, n= 10-13 slices, low gamma *p*=0.0004, high gamma *p*= 0.0026, one-way ANOVA with Sidak’s multiple comparisons) as is the difference between control and CORT in the low gamma range (p= 0.045, one-way ANOVA with Sidak’s multiple comparisons). In contrast the difference between controls and CORT +VFX is not significant. Shown in 5C are representative tracings from the 3 groups as indicated. As shown in figures 5D and E, venlafaxine’s effect in MMP-9 knockout animals was reduced as compared to its effect in the wild type. There is no significance difference between the CORT and CORT +VFX groups in these animals (4-5 mice, n= 9-11 slices, low gamma *p*= 0.96 high gamma *p*=0.63, one-way ANOVA with Sidak’s multiple comparisons). In 5F we show representative tracings from the knockout animals.

### 6. VFX stimulates an MMP-9-dependent increase in SWR abundance in ex vivo hippocampal slices from CORT-treated animals

While previous studies have linked major depressive disorder and animal models of the same to reduced gamma power, studies examining hippocampal sharp wave ripple (SWR) abundance in major depression or animal models are lacking. Rapid sequential replay of hippocampal neuronal assemblies, initially activated in the same sequence during learning, occurs during SWRs and is important to memory consolidation (Ego-Stengel and Wilson, 2010). Importantly, declarative memory consolidation is reduced in patients with MDD (Nissen et al., 2010). We therefore examined SWR abundance in control and corticosterone-treated animals. As shown in figure 6A, VFX increases SWR abundance in CORT-treated wild type animals (4-6 mice, n=9-13 slices, *p*=0.0003, one-way ANOVA with Tukey’s multiple comparisons). The difference between control and CORT + VFX was also significant *(p*= 0.049). Representative traces for wild type animals are shown in 6B. In contrast, as shown in figure 6C, VFX’s effect on SWR abundance in MMP-9 knockout animals was not significant (4-5 mice, n= 9-10 slices, *p*=0.308, one-way ANOVA with Tukey’s multiple comparisons). Shown in 6D are representative tracings for MMP-9 null animals.

**6.**
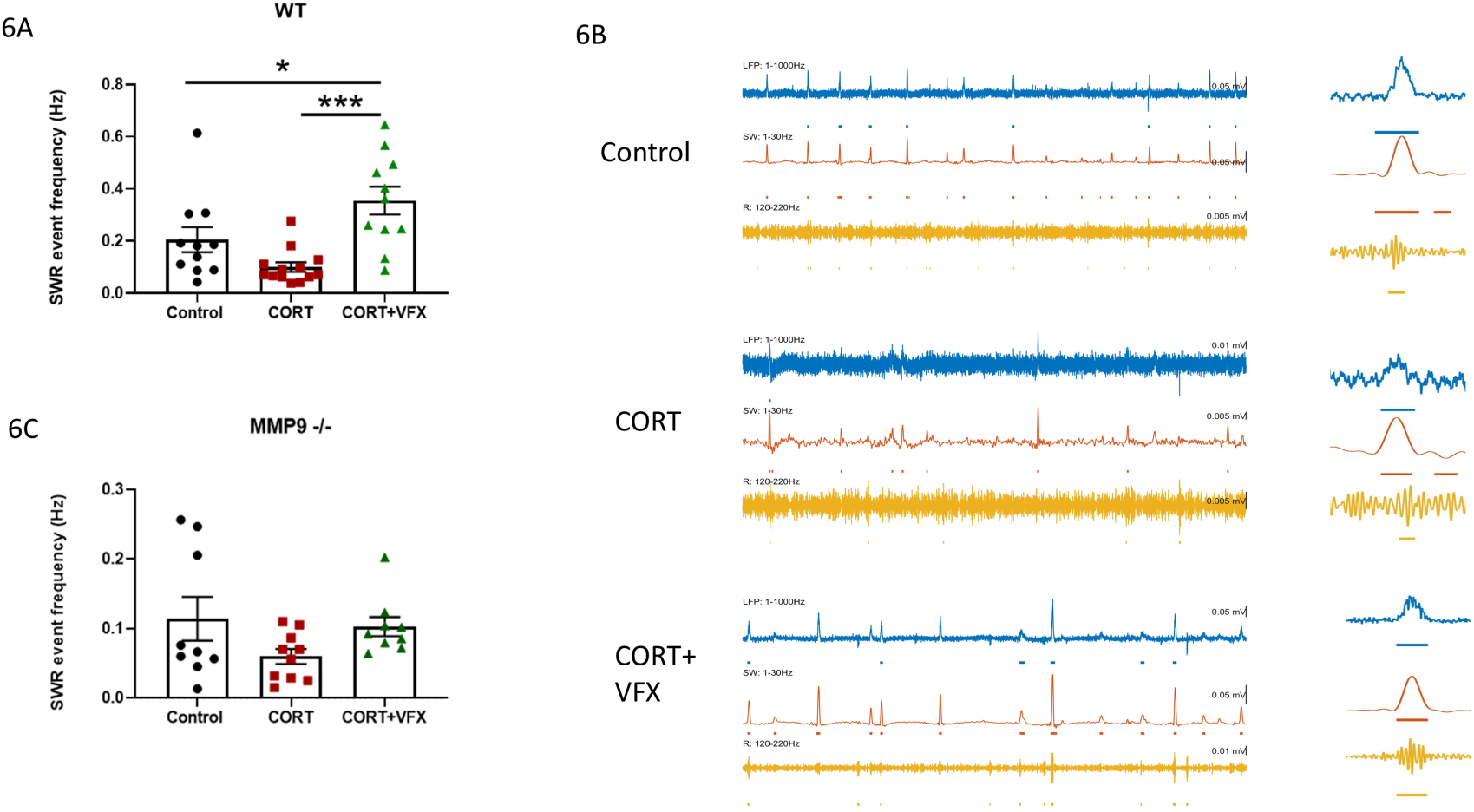
Venlafaxine stimulates an MMP-9-dependent increase in SWR abundance in ex vivo hippocampal slices from corticosterone treated animals. As shown in figure 6A, VFX increases SWR abundance in CORT-treated wild type animals (4-6 mice, n=9-13 slices, *p*=0.0003, one-way ANOVA with Tukey’s multiple comparisons). The difference between control and CORT + VFX was also significant *(p*= 0.049). Representative traces for wild type animals are shown in 6B. In contrast, as shown in figure 6C, VFX’s effect on SWR abundance in MMP-9 knockout animals was not significant (4-5 mice, n= 9-11 slices, *p*=0.308, one-way ANOVA with Tukey’s multiple comparisons). Representative traces for MMP-9 null animals are shown in 6D.

### 7. Increased anxiety and reduced working memory in CORT-treated mice; rescue by VFX in wild type mice

Corticosterone has been shown to increase anxiety as assessed by elevated plus maze (EPM) testing in C57BL/6 mice, with rescue by fluoxetine. Herein we examine the effect of VFX on this endpoint. In addition, since increased gamma power has been linked to enhanced working memory (Alekseichuk et al., 2016; Goodman et al., 2018; Martorell et al., 2019, Iaccarino et al., 2016), we also examine the effect of CORT and VFX on spontaneous alternations in the T maze, which has been shown to measure spatial working memory in mice (Shoji et al., 2012).

Consistent with a reduction in anxiety-like behavior, VFX reduced closed arm time in CORT-treated animals tested with the EPM (Figure 7A, n=7-9 mice per group, *p*= 0.029, one-way ANOVA with Sidak’s multiple comparisons testing). There was no significant difference between the CORT and CORT + VFX groups in MMP-9 null animals (Figure 7B, n=7-9 animals per group, *p*= 0.67, ANOVA with Sidak’s multiple comparisons testing). In Figures 7C, we examined the effects of CORT on T maze performance. CORT decreased working memory in wild type mice (n=10 mice per group, *p*= 0.0049, Student’s t-test). We also examined working memory in wild type and MMP-9 null mice treated with CORT or CORT +VFX (7C-D). VFX improved T-maze performance in CORT-treated control (n= 5 mice per group, *p* = 0.045, ANOVA with Sidak’s multiple comparisons testing). In addition, VFX improved performance in animals that did not receive CORT (n= 5 mice per group, *p* = 0.045, ANOVA with Sidak’s multiple comparisons testing). The difference between control and CORT groups was not significant in this cohort, which may be due to injection stress in both groups and/or a smaller n. In addition, though the wild type controls in 7C and 7D showed differential performance, it should be noted that controls in figure 7D received stressful daily intraperitoneal saline while controls in figure 7C received β-CD in their drinking water. In Figure 7E, we show T maze results for MMP-9 null animals. We did not have a VFX only group for these animals. However, in contrast to wild type animals, VFX did not improve performance in CORT-treated MMP-9 null mice (n=6-7 animals per group, *p*= 0.82, ANOVA with Sidak’s multiple comparisons testing).

**7.**
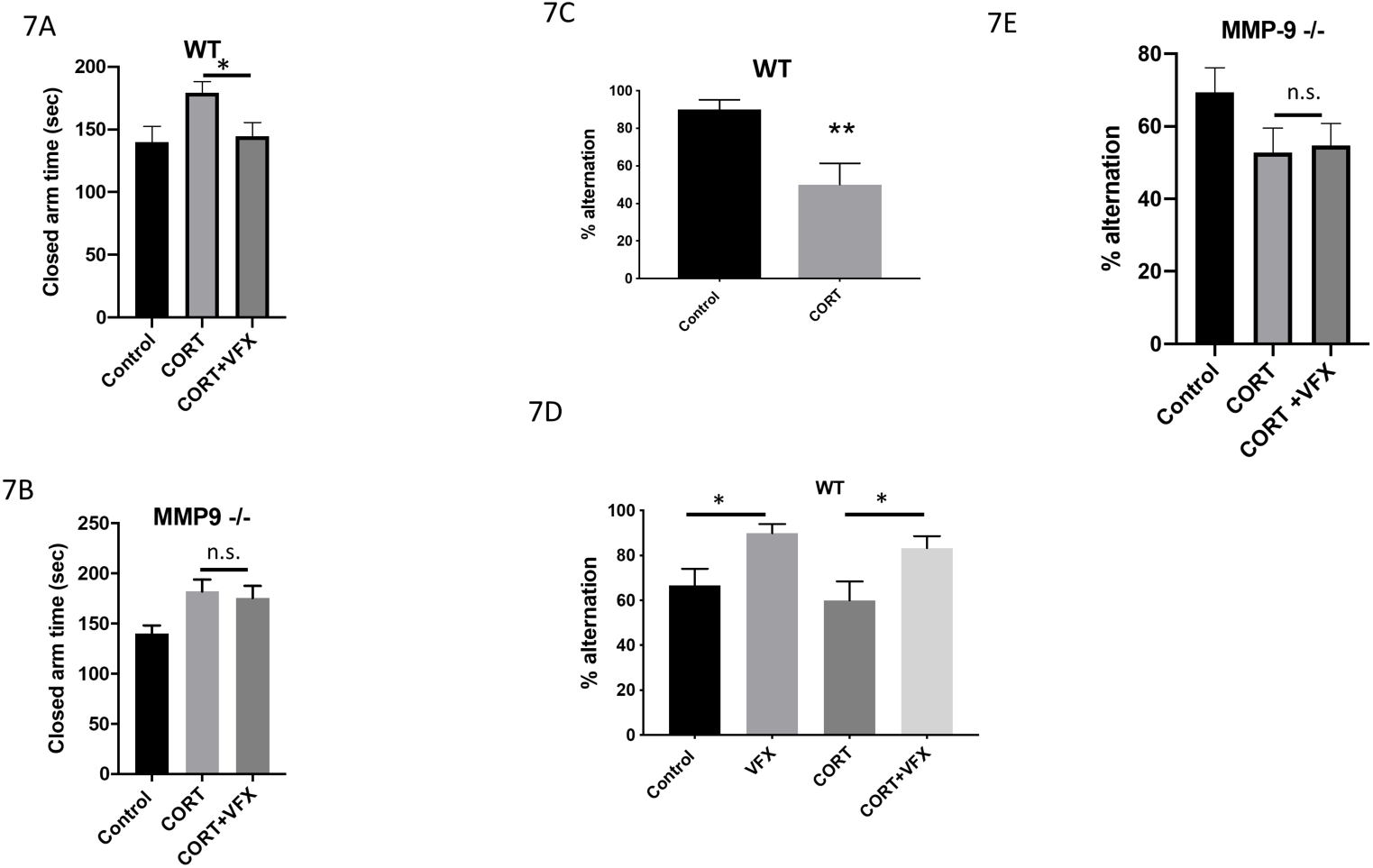
Reduced working memory in corticosterone-treated mice; rescue by VFX in wild type mice. As shown in figure 7A, VFX reduced closed arm time in CORT-treated animals tested with the EPM (n=7-9 mice per group, *p*= 0.029, one-way ANOVA with Sidak’s multiple comparisons testing). In contrast, there was no significant difference between the CORT and CORT + VFX groups in MMP-9 null animals (Figure 7B, n=7-9 animals per group, *p*= 0.67, ANOVA with Sidak’s multiple comparisons testing). In Figures 7C, we examined the effects of CORT on T maze performance. CORT decreased working memory in wild type mice (n=10 mice per group, *p*= 0.0049, Student’s t-test). We also examined working memory in wild type and MMP-9 null mice treated with CORT or CORT +VFX (7C-D). VFX improved T-maze performance in CORT-treated control (n= 5 mice per group, *p* = 0.045, ANOVA with Sidak’s multiple comparisons testing). In addition, VFX improved performance in animals that did not receive CORT (n= 5 mice per group, *p* = 0.045, ANOVA with Sidak’s multiple comparisons testing). The difference between control and CORT groups was not significant in this cohort, which may be due to injection stress in both groups and/or a smaller n. In addition, though the wild type controls in 7C and 7D showed differential performance, it should be noted that controls in figure 7D received stressful daily intraperitoneal saline while controls in figure 7C received β-CD in their drinking water. In Figure 7E, we show T maze results for MMP-9 null animals. We did not have a VFX only group for these animals. However, in contrast to wild type animals, VFX did not improve performance in CORT-treated MMP-9 null mice (n=6-7 animals per group, *p*= 0.82, ANOVA with Sidak’s multiple comparisons testing).

### 8. MMP-9/TIMP-1 levels are increased in PFC samples from MDD patients treated with monoamine antidepressants

PNN alterations have been examined in depression and animal models of the same. Sulfation patterns that render nets more resistant to proteolysis have been observed in human postmortem tissue from depressed patients and increased PNN deposition has been noted in animal models of both stress and depression (Pantazopoulos et al., 2015; Riga et al., 2017). Brain levels of PNN modulating proteases have, however, not been well examined. To assess the possibility that MMP-9 levels may be altered with depression and/or increased by monoamine antidepressants, we analyzed post-mortem PFC lysates from control patients, untreated major depressive disorder patients, and monoamine antidepressant-treated major depressive disorder patients for MMP-9 protein based on an ELISA. We also analyzed these lysates for levels of tissue inhibitor of metalloproteinase-1, an endogenous inhibitor of MMP-9. Results are shown in figures 8A and B, and demonstrate that as compared to control patients, monoamine antidepressant-treated patients (anti-depressant drug or ADD) have increased levels of MMP-9 in the prefrontal cortex (8A, *p* = 0.0130, one-way ANOVA with Dunnett’s multiple comparisons, 2 high outliers in the MDD/ADD group were excluded). In contrast, there is no significant difference between MMP-9 levels in control and untreated MDD patients (*p*=0.45). Moreover, there is a significant increase in the MMP-9/TIMP-1 ratio in monoamine antidepressant-treated MDD patients as compared to the same ratio in controls or untreated MDD patients (Control versus MDD + ADD, *p*= 0.0052, one-way ANOVA with Tukey’s multiple comparisons; MDD versus MDD + ADD, *p*=0.009, one-way ANOVA with Tukey’s multiple comparisons).

**8.**
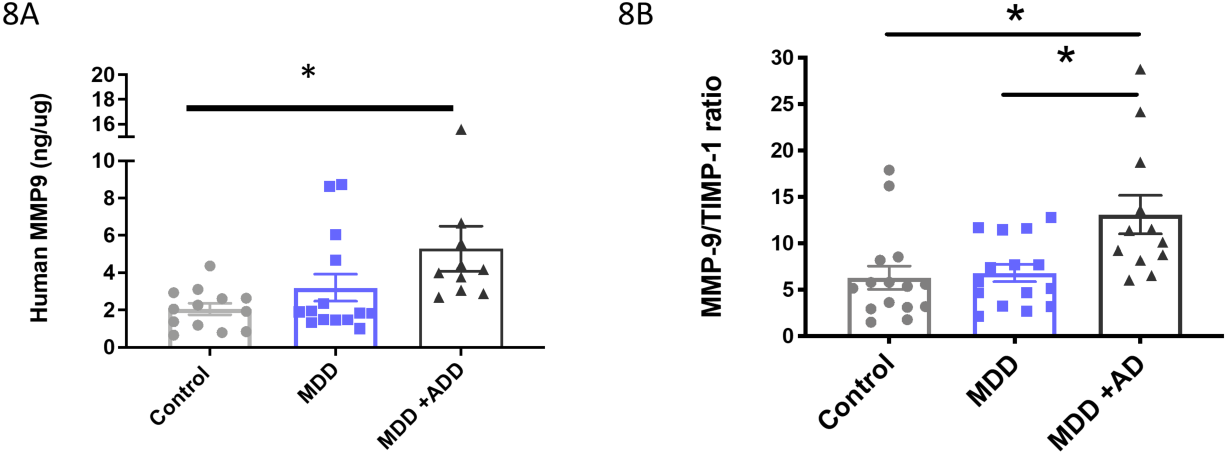
MMP-9/TIMP-1 levels are increased in PFC samples from MDD patients treated with monoamine antidepressants. Results shown in figures 8A and B demonstrate that as compared to control patients, monoamine antidepressant-treated patients (anti-depressant drug or ADD) have increased levels of MMP-9 in the prefrontal cortex (8A, *p* = 0.0130, one-way ANOVA with Dunnett’s multiple comparisons). In contrast, there is no significant difference between MMP-9 levels in control and untreated MDD patients (*p*=0.45). Moreover, there is a significant increase in the MMP-9/TIMP-1 ratio in monoamine antidepressant-treated MDD patients as compared to the same ratio in controls or untreated MDD patients (Control versus MDD + ADD, *p*= 0.0052, one-way ANOVA with Tukey’s multiple comparisons; MDD versus MDD + ADD, *p*=0.009, one-way ANOVA with Tukey’s multiple comparisons).

**9.**
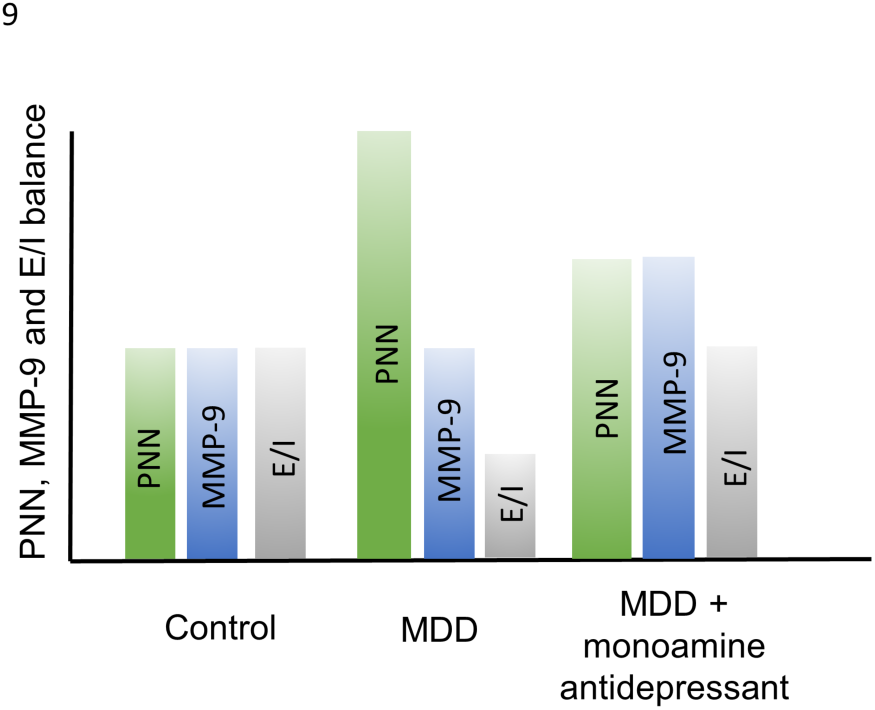
Hypothetical changes in PNN deposition and MMP-9 activity in untreated and treated MDD. Hypothetical PNN, MMP levels, and E/I balance (arbitrary units) are shown for non-depressed control patients. With MDD, PNN levels increase and thus E/I balance decreases. With MDD in the setting of antidepressant therapy, however, MMP levels also increase to potentially restore E/I balance.

Of note, there was no significant difference in MMP-9/TIMP-1 ratio as a function of sex (n=24 males and 17 females, average MMP-9/TIMP-1 in males 7.82 +/- 0.87 S.E.M., average MMP-9/TIMP-1 in females 8.34 +/- 1.68 S.E.M.). Moreover, in post-unblinding exploratory analyses, limited by a low n due to cause of death ambiguity for some patients, we compared MMP-9/TIMP-1 ratios in MDD patients who committed suicide versus those who did not. The average MMP-9/TIMP-1 in suicide patients (n=9) was 1.64 +/- 1.3 S.E.M. and the ratio in non-suicide patients (n=4) was 7.02 +/- 0.99 S.E.M.

### 9. Schematic; hypothetical changes in PNN deposition and MMP-9 activity in untreated and treated MDD

Stress and depression can enhance PNN deposition and also render PNNs resistant to proteolysis (Pantazopoulos et al., 2015; Riga et al., 2017; Simard et al., 2018). Both processes can increase overall PNN levels. Monoamine reuptake inhibitors can, however, act to increase expression of PNN degrading proteases and normalize PNN levels. Though excess PNN remodeling may enhance neuronal plasticity at the cost of memory stability (Gogolla et al., 2009; Slaker et al., 2015; Carulli et al., 2016), moderately increased PNN remodeling in the background of excess deposition may be adaptive or therapeutic. Stress might also reduce dendritic arbor, through PNN dependent and/or independent mechanisms, and this may be normalized by monoamine antidepressant associated changes in MMP-9 (not depicted).

## Discussion

Herein, we tested the hypothesis that VFX stimulates MMP-9 dependent changes relevant to increased E/I balance in conventionally housed and/or corticosterone treated mice. We also tested an association between antidepressant use and MMP-9 levels in humans. Our hypothesis was based on prior work linking MMP-9 to endpoints including enhanced hippocampal-dependent learning and memory (Nagy et al., 2006), as well as monoamine-stimulated MMP-9 activity (Bijata et al., 2017).

In work related to monoamines and MMP-9 release, a 5HT7 agonist stimulates MMP-9-dependent growth of dendritic spines in hippocampal cultures and slices (Bijata et al., 2017). 5HT7 is coupled to Gα_s_, which increases cAMP levels and cfos-dependent transcription (Boutillier et al., 1992). The latter is a critical transcription factor for MMP-9 expression (Kaczmarek, 2018). Serotonin reuptake inhibitors are also thought to act on Gα_s_ coupled receptors to increase α secretase activity (Fisher et al., 2016). With ADAM-10, MMP-9 represents a potent α secretase (Fragkouli et al., 2012). Because VFX can increase both serotonin and norepinephrine signaling, we examined MMP-9 release from cultured hippocampal neurons stimulated with β adrenergic agonists that activate Gα_s_ coupled receptors. Isoproterenol indeed increased neuronal MMP-9 release.

In work more directly linked to our central hypotheses, VFX stimulated MMP-9-dependent increases in cortical dendritic arbor and hippocampal PSD-95 expression. Consistent with this, previous studies have shown that MMP-9 can induce spine head protrusions (Szepesi et al., 2013) in organotypic cultures, and spine expansion in hippocampal slices (Wang et al., 2008). Moreover, a pan-MMP inhibitor reduces neurite outgrowth in cultured cortical neurons (Ould-yahoui et al., 2009).

We acknowledge that the MMP-9 dependence of our findings is inferred from comparison to constitutive knockouts. Though alternatives include the use of MMP inhibitors, these inhibitors are non-specific (Fields, 2015). In the absence of inducible MMP-9 knockouts, constitutive knockouts have demonstrated MMP-9-dependent LTP and MMP-9-dependent deficits in a mouse model of Fragile X syndrome (Nagy et al., 2006; Wen et al., 2018). Constitutive knockouts have also demonstrated MMP-9-dependent plasticity following light reintroduction after dark (Murase et al., 2017). Consistent with prior studies (Tian et al., 2007), baseline differences in PSD-95 between MMP-9 knockouts and controls were not observed in adult animals. Baseline differences in somatosensory dendritic arbor could not be compared, since wild type animals with and without VFX were stained at one time and knockouts at another. Dendritic arbor results were, however, normalized to the appropriate control.

We also observed an MMP-9-dependent reduction in overall PNN intensity and PV/PNN ratios in wild-type animals. MMP-9 knockouts also demonstrated reduced basal and VFX-stimulated cleavage of BCAN while levels of full length BCAN were not reduced in MMP-9 knockouts. One possibility is that BCAN turnover is increased in wild-type as compared to MMP-9 null mice. Specifics of PNN composition and/or turnover may be functionally relevant (Carulli et al., 2010; Sorg et al., 2016). Cleaved BCAN may function differently than full length, and newly synthesized BCAN could be poorly localized.

The MMP-9 dependence of basal and VFX-associated changes in BCAN cleavage is interesting given that PNN enwrapped PV neurons are not noted to express MMP-9 under basal conditions (Rossier et al., 2015). MMP-9 could be released from excitatory inputs to PV expressing neurons. Indeed, MMPs have been noted at synaptic boutons (Shilts and Broadie, 2017).

We additionally observed that BCAN levels and PNN staining were increased in the corticosterone model of depression, with the latter normalized by VFX in wild-type mice. These results are consistent with those from a study in a social defeat stress in rats, in which levels of PNN components were increased (Riga et al., 2017). Imipramine normalized stress-associated changes in PNN, though underlying molecular mechanisms were not examined (Riga et al., 2017). Since PNN reductions are linked to pyramidal cell disinhibition (Hayani et al., 2018), our work supports a link between monoamine reuptake inhibition and increased in E/I balance.

We thus tested E/I balance in *ex vivo* murine slices using an approach based on neuronal population activity. Population recordings have the advantage of detecting relatively broad changes in E/I balance, and may also detect alterations if PNN disruption is limited to a percentage of cells. Changes in single cell recordings may instead depend on whether recordings are made in specific cells in which PNN changes are appreciable. Consistent with the possibility that increased PNN deposition enhances PV interneuron-mediated inhibition, CORT treatment reduced SWR abundance and gamma power in *ex vivo* slices in wild-type mice. Moreover, VFX stimulated an MMP-9-dependent increase in SWR abundance and carbachol-stimulated gamma power in *ex vivo* hippocampal slices from CORT-treated wild-type but not MMP-9 null animals. Similar to other neurophysiological endpoints (Nagy et al., 2006; Ferrer-Ferrer and Dityatev, 2018), significant basal differences in gamma power were not observed.

Gamma power represents a physiological correlate of increased E/I balance and is critical to memory (Zheng et al., 2016; Lensjo et al., 2017). Increased hippocampal gamma power improves pattern completion and the quality of retrieved spatial memories (Staresina et al., 2016; Stevenson et al., 2018). Increases in theta and gamma power during encoding also predict recall (Sederberg et al., 2003). Gamma oscillations are critical to working memory (Yamamoto et al., 2014), which may be limited by the number of gamma cycles nested within a theta cycle (Lisman, 2010). VFX’s enhancement of gamma power is relevant to depression since gamma power is altered in depression (Liu et al., 2012) and increased with remission in both humans and animal models (Khalid et al., 2016; Arikan et al., 2018). Moreover, increased gamma power correlates with improvements on the Hamilton depression scale (Arikan et al., 2018).

VFX’s enhancement of SWR abundance is also relevant to depression. During SWRs, cell assemblies that were sequentially activated during encoding are sequentially reactivated in a time-compressed manner during off-line states. SWRs are critical to memory consolidation (Ego-Stengel and Wilson, 2010; Buzsaki, 2015), which may be impaired in MDD (MacQueen and Frodl, 2011). Further study on the quality of SWRs in terms of reactivation of learning-relevant assemblies, as opposed to random assemblies, are warranted.

Since PV fast spiking interneuron activity is critical to gamma and ripple expression, it is of interest that despite reduced PV activity with PNN attenuation (Hayani et al., 2018; Tewari et al., 2018), gamma oscillations and SWR events are not only maintained but increased in power and event frequency. This is consistent with the possibility that the PV population retains responsivity when excitatory input is strong, and is supported by previous work (Balmer, 2016).

A caveat is that MMP-9 has substrates in addition to PNN components that may increase pyramidal cell activity and thus gamma power or SWR abundance through relatively direct effects on excitatory neurons. These mechanisms include rapid MMP-dependent generation of integrin-binding ligands and integrin-stimulated expansion of spines on glutamatergic neurons (Lonskaya et al., 2013). In addition, a small population of pyramidal neurons in CA2 hippocampus are PNN-enveloped, which may restrict their ability to undergo LTP (Carstens et al., 2016). However, given the abundance of the PV neuron-associated PNN and the potential for each PV+ cell to impact a number of pyramidal cells (Andersen et al., 2006), we posit that disruption of PV neuron-associated PNNs makes a strong contribution to these endpoints in or study. Future studies with mutants that render nets relatively resistant to proteolysis may be useful for additional support (Foscarin et al., 2017).

Previous work has examined effects of monoamine antidepressants on long-term memory (Van Dyke et al., 2019). Because of the link between gamma power and working memory, behavioral studies herein focused on short-term memory. T-maze performance was reduced with corticosterone and enhanced by VFX in conventionally housed and corticosterone-treated mice. In contrast, VFX did not enhance working memory in MMP-9 null mice. These results are consistent with the effects of the PNN-degrading enzymes on object place recognition (Riga et al., 2017) and reversal learning (Happel et al., 2014).

Consistent with the possibility that monoamine antidepressants stimulate MMP-9 expression in humans, we observed increased levels of MMP-9 and increased MMP-9/TIMP-1 ratios in antidepressant-treated individuals. TIMP-1 is inducible and expressed in areas relevant to depression. TIMP-1 inhibits MMP-9 to a greater degree than does TIMP-2 and moreover, TIMP-1 abolishes MMP-9-dependent LTP in prefrontal cortex (Okulski et al., 2007). Our initial hypothesis was that MMP-9 levels would be lower in the depressed group and normalized by antidepressant treatment. The observation that treated patients have higher MMP-9/TIMP-1 ratios than control or depressed individuals suggests that changes occurring with depression, such as increased PNN deposition, may instead be normalized by treatment-associated increases in MMP-9.

In conclusion, our study demonstrates that VFX stimulates MMP-9-dependent increases in dendritic arbor and PSD-95 expression. We are also the first to show MMP-9 dependent changes in *ex vivo* gamma power, SWR abundance and working memory in a corticosterone model of depression. Future studies may address the question of whether these changes are largely beneficial and whether agents that target PNN integrity and/or gamma power, such as GENUS (Martorell et al., 2019), might also be useful for the treatment of depression.

## Acknowledgements

We would like to acknowledge outstanding veterinary support from the Department of Comparative Medicine and we would also like to apologize to investigators whose excellent work was not directly cited. We would also like to thank Adam Caccavano for software development and sharing and assistance with images. Katherine Conant received funds for support and supplies from NIMH (R21MH118749) as well as Deborah Wilson and Anthony Herman through the Georgetown University *Partners in Research* program. Seham Alaiyed received support from the Saudi Arabian government, Qassim University scholarship program. Mondona McCann was supported with funding from T32 NS04121. The authors deeply appreciate the invaluable contributions made by the families consenting to donate brain tissue and be interviewed. We also gratefully acknowledge the support of the staff of the Cuyahoga County Medical Examiner’s Office, Cleveland, Ohio. We gratefully acknowledge the assistance of Drs. James Overholser and George Jurjus, and of Lesa Dieter in the psychiatric assessments, and of Timothy M. De Jong and Lisa Konick in acquiring written consent and collecting tissues. The work performed by the Postmortem Brain Core is supported, in part, by funds from the IDeA Program of the National Institute of General Medical Sciences of the National Institutes of Health, Center for Psychiatric Neuroscience (CPN)-COBRE (P30GM103328) and by National Institute of Mental Health (R01 MH67996). There are no competing financial interests for any of the authors in relation to the work described.

